# Structural Analysis of the PATZ1 BTB domain homodimer

**DOI:** 10.1101/2020.01.13.903898

**Authors:** Sofia Piepoli, Aaron Oliver Alt, Canan Atilgan, Erika J. Mancini, Batu Erman

## Abstract

PATZ1 is a transcriptional repressor belonging to the ZBTB family that is functionally expressed in T-lymphocytes, as well as in a ubiquitous fashion. PATZ1 targets the *Cd8* gene in lymphocyte development and interacts with the p53 protein to control genes important in proliferation and DNA damage response. PATZ1 exerts its activity through an N-terminal BTB domain that mediates dimerization and co-repressor interactions and a C-terminal zinc finger motif-containing domain that mediates DNA binding. Here, the crystal structures of the murine and zebrafish PATZ1 BTB domains are reported at 2.3 and 1.8 Å resolution respectively. The structures reveal that, like other ZBTB structures, the PATZ1 BTB domain forms a stable homodimer and likely binds co-repressors through a lateral surface groove that is formed upon dimerization. Analysis of the lateral groove reveals a large acidic patch in this region which contrasts previously resolved basic co-repressor binding interfaces in other ZBTB proteins. A large 30 amino acids glycine- and alanine-rich central loop, unique to mammalian PATZ1 amongst all ZBTB proteins, could not be resolved likely due to its flexibility. Modelling of this loop indicates that it can participate in the dimerization interface of BTB monomers.

**Synopsis:** The crystal structures of the PATZ1 BTB domain in mammals and fish are homodimers. The core dimer conformation of these BTB proteins is dynamically stable, despite the presence of highly flexible regions in the dimerization interface.

## 1. Introduction

PATZ1 (POZ-, AT hook-, and Zinc finger-containing protein 1) also known as ZBTB19, is a transcription factor present in all vertebrates (Fig. 1A). It was first discovered in a Yeast Two Hybrid (Y2H) experiment in which it associated through its BTB (Broad-Complex, Tramtrack and Bric a brac) domain with the BTB domain of the transcription factor BACH2 (BTB And CNC Homology 2) (Kobayashi *et al.*, 2000). PATZ1 is also referred to as MAZR (Myc-Associated Zinc finger Related) because of the close similarity between its zinc finger (ZF) domain and that of MAZ (Myc-Associated Zinc finger). While its expression can be detected in many cell types and developmental stages, PATZ1/ ZBTB19/ MAZR is highly expressed specifically in the early stages of T-lymphocyte differentiation where it negatively regulates *Cd8* gene expression (Bilic & Ellmeier, 2007). PATZ1 has been shown to participate in thymocyte development and CD4, CD8 and T_reg_ lineage choice by repressing the expression of ThPOK (ZBTB7B/ ZBTB15/ cKrox), another BTB domain-containing transcription factor (Sakaguchi et al., 2010, Sakaguchi et al., 2015, He et al., 2010) and the expression of the FOXP3 transcription factor (Andersen *et al.*, 2019). The functions of PATZ1 are however not limited to lymphocytes as its expression is ubiquitous. An early embryonic role for PATZ1 has been suggested, as Patz1-/- mice are embryonic lethal or born at non-Mendelian frequency and are small in size depending on the genetic background (Sakaguchi et al., 2010). PATZ1 also negatively regulates induced pluripotent stem cell (iPSC) generation (Ow *et al.*, 2014, Ma *et al.*, 2014). This function may be related to its interaction with the p53 tumor suppressor as demonstrated by various studies (Valentino, Palmieri, Vitiello, Pierantoni, *et al.*, 2013, Keskin *et al.*, 2015, Chiappetta *et al.*, 2015).

**Figure 1.**
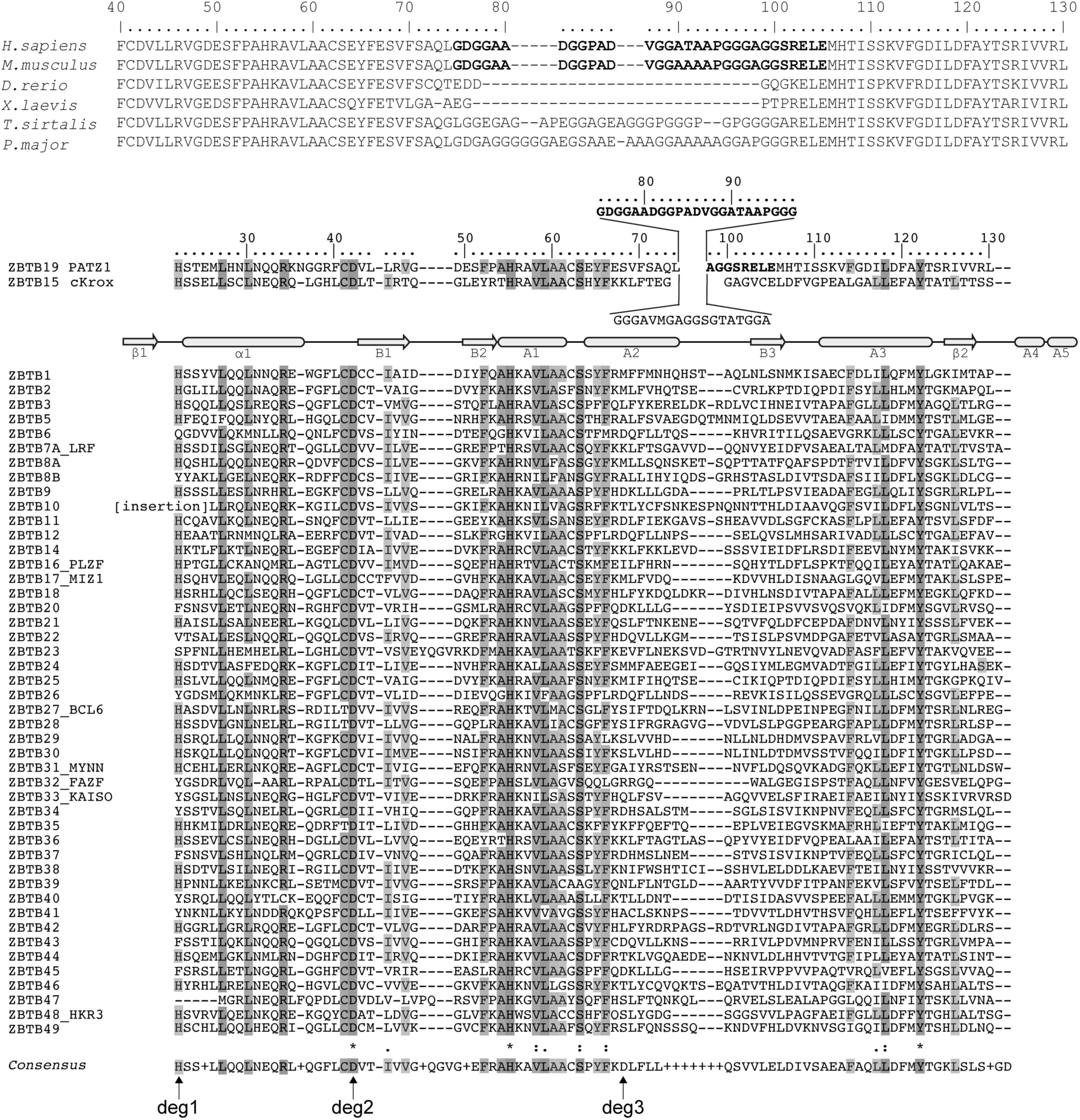
Sequence alignment of BTB domains of ZBTB transcription factors identifies a unique central region in PATZ1, conserved in mammals. **(A)** Sequence alignment of PATZ1 BTB domains from selected vertebrate species. Unique central sequence of the A2/B3 loop is conserved in mammals and missing in fish and amphibians is indicated in bold letters. **(B)** Sequence alignment of selected human ZBTB proteins and their predicted secondary structure. The sequence of the human PATZ1 and cKrox BTB domains with their unique extra region between the A2 helix and B3 strand are shown above. PATZ1 amino acids that correspond to this region, without electron density assignments from the crystal structure are in bold letters. Arrows and rods identify predicted conserved β-strand and α-helical regions. The eight BTB domains with solved structures are annotated on the left with their common names in addition to the ZBTB nomenclature. Shading, asterisk, column and period symbols identify conserved residues according to the Clustal format. A consensus sequence is shown at the bottom with three predicted degron residues involved in BTB domain stability indicated by arrows.

Structurally, PATZ1 belongs to the POZ (Pox virus and Zinc finger) or ZBTB (Zinc finger and BTB) family of transcription factors (Lee & Maeda, 2012). Proteins belonging to this family have been implicated in many biological processes, including transcriptional regulation and development (Chevrier & Corcoran, 2014) whilst their dysfunction in vertebrates has been linked to tumorigenesis. ZBTB proteins bind to DNA through their ZF domains and use their BTB domains for oligomerization (Bonchuk *et al.*, 2011) and recruitment of co-repressors and chromatin remodelling factors (Bardwell & Treisman, 1994, Siggs & Beutler, 2012). The human genome encodes for 49 members of the ZBTB family (Fig. 1B), all of which contain an N-terminal BTB domain and a variable number of ZF motifs at their C-terminus. Two members of this family contain an additional motif in the form of an AT hook (PATZ1 and PATZ2/ ZBTB24).

The BTB domain is a structural feature that mediates functional interactions between proteins (Perez-Torrado et al., 2006). Most of the available BTB domain structures of the ZBTB family are homodimers formed by the assembly of two identical monomers (Ahmad et al., 1998, Li et al., 1999, Ahmad et al., 2003, Schubot et al., 2006, Stogios et al., 2007, Ghetu et al., 2008, Stead et al., 2008, Cerchietti et al., 2010, Stogios et al., 2010, Sakamoto et al., 2017, McCoull et al., 2017, Kamada et al., 2017, Yasui et al., 2017, Kerres et al., 2017, McCoull et al., 2018, Cheng et al., 2018). In the case of Myc-interacting zinc finger protein 1 (MIZ1), the structures of both a homodimer (Stogios et al., 2010) and a homo-tetramer (Stead et al., 2007) have been reported.

The overall fold of the BTB domain is highly conserved, containing the following secondary structure elements: β1-α1-B1-B2-A1-A2-B3-A3-β2-A4-A5 (Stogios et al., 2005). A unique feature of the PATZ1 BTB domain is a long glycine- and alanine-rich central loop between A2 and B3 (Kobayashi et al., 2000) (Fig. 1). This central loop is conserved in all vertebrate PATZ1 proteins, but it is absent in fish and amphibians. The two BTB monomers are known to homo-dimerize through a specific dimerization interface that includes β1-α1; A1-A2; β2-A5. Although homodimerization seems to be favored, heterodimeric interactions between couples of ZBTB proteins have also been documented. The crystal structure of a tethered “forced” heterodimer between the BTB domains of MIZ1 and B-cell lymphoma 6 protein (BCL6) suggests that heterodimers can use the same interface as homodimers (Stead & Wright, 2014). Together with BCL6, MIZ1 seems to be a promiscuous member of the ZBTB family, making more heterodimers than any other BTB domain. MIZ1 functionally interacts with BCL6 in germinal center B cells (Phan et al., 2005), ZBTB4 (Weber et al., 2008) and NAC1 (nucleus accumbens-associated-1; (Stead & Wright, 2014)). PATZ1 can also form heterodimers with other BTB domain containing proteins as with BCL6, PATZ2 (Huttlin et al., 2015), BACH1 and BACH2 (Kobayashi et al., 2000).

The mechanism controlling homodimerization versus heterodimerization of BTB domains has not been elucidated. Co-translational dimerization, a mechanism often required in protein complex assembly, may be at play (Kramer et al., 2019). Recently, a dimerization quality control mechanism for BTB proteins was proposed, where homodimer stability would exceed that of heterodimers because of the structural masking of destabilizing residues (Herhaus & Dikic, 2018, Mena et al., 2018). According to this model, the preferential exposure of three “degron” residues on BTB heterodimers results in targeting by ubiquitin ligases and a shorter half-life. Whether this mechanism is universally shared by all BTB domain-containing proteins including members of the ZBTB family remains unclear.

Homodimerization of BCL6 and Promyelocytic leukemia zinc finger (PLZF) proteins creates a charged groove that binds nuclear receptor co-repressors such as NCOR1, NCOR2 (SMRT) and BCOR (Huynh & Bardwell, 1998, Wong & Privalsky, 1998, Huynh *et al.*, 2000, Melnick *et al.*, 2000, Melnick *et al.*, 2002). NCOR1 and SMRT are structurally disordered proteins that share 45% identity (Granadino-Roldan et al., 2014) and contain a conserved 17-amino acids long BCL6 Binding Domain (BBD). These co-repressors are components of large complexes containing histone deacetylases (Li et al., 2000) contributing to transcriptional silencing. It is not known if co-repressor binding is a generalizable feature of BTB homodimers, as MIZ1, FAZF and LRF BTB homodimers do not interact with these co-repressors (Stogios *et al.*, 2010, Stogios *et al.*, 2007). The PATZ1 BTB domain has been shown to interact with NCOR1, however it is not known if the interaction is mediated by an interface similar to that of BCL6 and PLZF (Bilic et al., 2006).

Before this work, the structures of eight different proteins belonging to the ZBTB family were available in the Protein Data Bank (PDB): LRF/ Pokémon (ZBTB7A); PLZF (ZBTB16); MIZ1 (ZBTB17); BCL6 (ZBTB27); MYNN (ZBTB31); FAZF (ZBTB32); KAISO (ZBTB33); HKR3/ TZAP (ZBTB48). In order to obtain biological insights into the binding of PATZ1 to co-repressors we set off to determine the atomic structure of the PATZ1 (ZBTB19) BTB domain. To probe the role of the unique A2/B3 central loop, we obtained the crystal structures of both the mouse PATZ1 BTB domain and of its zebrafish orthologue.

## 2. Materials and methods

### 2.1. Protein expression and purification

The human isoform 3 (Q9HBE1-3) and the mouse PATZ1 BTB (Q9JMG9) sequences are 98.9% identical (LALIGN (Gasteiger et al., 2003)), diverging only for one residue (T91A). The mouse PATZ1 BTB protein coding sequence (12-166) was PCR-amplified from a CMV-HA plasmid construct (Keskin et al., 2015) containing the full-length mouse PATZ1 cDNA, cloned into a pET-47b bacterial expression plasmid (Novagen) between SmaI and NotI restriction sites in frame with an N-terminal 6x His-tag and an HRV 3C Protease cleavage site for fusion tag removal. The resulting plasmid was transformed into the *E. coli* Rosetta-2 DE3 strain and grown at 310 K by shaking at 180 rpm in Terrific/ Turbo Broth (TB) medium supplemented with 50 µg/ml Kanamycin and 33 µg/ml Chloramphenicol until the absorbance at 600 nm reached a value of 0.6. Expression of the fusion protein was induced by the addition of 0.1 mM isopropyl β-D-1-thiogalactopyranoside (IPTG) and growth was continued for 16 hours at 291 K. The zebrafish (*Danio rerio*) PATZ1 BTB protein coding sequence (1-135) was PCR-amplified from zebrafish genomic DNA (kind gift of Dr. S.H. Fuss), cloned and expressed in the same bacterial expression plasmid as described above.

The cells were harvested by centrifugation, resuspended in 25 ml of Lysis Buffer (50 mM HEPES pH 7, 250 mM NaCl, 10 mM imidazole, 0.5 mM TCEP, DNase, protease inhibitor cocktail) and disrupted by sonication on ice. The lysate was clarified by centrifugation at 26700 (x g) for 45 min at 277 K. The supernatant was applied onto a HisPur™ Cobalt Resin (ThermoFisher) column previously equilibrated with Wash Buffer (50 mM HEPES, 250 mM NaCl, 0.5 mM TCEP, 10 mM imidazole). Following 10 min incubation at 227 K and the application of Wash Buffer, the protein was then eluted by the addition of Elution Buffer (50 mM HEPES, 250 mM NaCl, 0.5 mM TCEP, 300 mM imidazole). The collected eluate was concentrated to 2.6 mg/ml using a Sartorius VivaSpin20 column (10 K MW cut-off) for additional purification by Size-Exclusion Chromatography (SEC) using a HiLoad^®^ 16/600 Superdex^®^ 75 prep grade column (GE Healthcare) in Gel Filtration Buffer (20 mM HEPES, 250 mM NaCl, 0.5 mM TCEP) at 277 K. Fractions were analyzed on a 14% SDS PAGE gel by electrophoresis and those containing PATZ1 BTB were pooled and concentrated to 9 mg/ml. The zebrafish PATZ1 BTB domain (69% sequence identity to mouse BTB) was expressed and purified as described above for the mouse BTB domain. The final concentration of the protein was 8.5 mg/ml.

### 2.2. Crystallization

All crystallization experiments were performed at 291 K using the sitting-drop vapor-diffusion method. Initial screening of 768 conditions was performed at the University of Sussex crystallization facility using a Crystal Phoenix dispensing robot to pipette 100 µl of protein solution and 100 µl of precipitant solution into single drops in 96-well plates. Crystals of the mouse PATZ1 BTB domain appeared after 72 hours in MMT (DL-Malic acid, MES monohydrate, Tris), 0.1 M PEG 1500 25% w/v. Crystals of the zebrafish PATZ1 BTB domain appeared after 72 hours in 40% v/v PEG 500 MME, 20% w/v PEG 20 K, 0.1 M Tris pH 8.5, 0.06 M Magnesium chloride hexahydrate, 0.06 M Calcium chloride dihydrate.

### 2.3. Data collection and processing

For data collection, single crystals were briefly immersed in 20% glycerol prior to flash-cooling in liquid nitrogen. For the mouse PATZ1 BTB, data to 2.29 Å were collected on Beamline I04 of the Diamond Light Source (Didcot, UK). Diffraction data was indexed, integrated, scaled, and reduced with XIA2 (Winter, 2010) and AIMLESS (Evans & Murshudov, 2013). The space group was P41212 (unit cell parameters a =b= 43.23, c = 162.95 Å, and α = β = γ = 90°) with one molecule in the asymmetric unit.

For the zebrafish PATZ1 BTB, data to 1.8 Å were collected on Beamline I04 of the Diamond Light Source (Didcot, UK). Diffraction data was indexed, integrated, scaled, and reduced with XIA2 and XDS (Kabsch, 2010). The space group was P3121 (unit cell parameters a = b= 43.08, c = 123.64 Å, and α = β = γ = 90°) with one molecule in the asymmetric unit.

Detailed X-ray data collection are given in Table 1.

**Table 1.** Data collection and refinement statistics.

|  | PATZ1 BTB Mouse | PATZ1 BTB Zebrafish |
| --- | --- | --- |
| Data Collection Statistics: |  |  |
| Wavelength (Å) | 0.97625 | 0.9795 |
| Space group | P 41 21 2 | P 31 21 |
| Cell dimensions: |  |  |
| a, b, c (Å) | 43.23, 43.23, 162.95 | 43.08, 43.08, 123.64 |
| $\alpha, \beta, \gamma$ (°) | 90, 90, 90 | 90, 90, 120 |
| Resolution (Å) | 43.23 - 2.29 (2.37 - 2.29) | 41.8 - 1.8 (1.9 - 1.8) |
| Number of unique reflections | 7573 (724) | 12786 (1120) |
| $R_{\text{merge}}$ | 0.05 (0.67) | 0.01 (0.49) |
| $I/\sigma I$ | 9.91 (1.39) | 19.96 (1.27) |
| Completeness (%) | 99.58 (99.18) | 98.41 (87.99) |
| Redundancy | 2.0 (2.0) | 2.0 (1.8) |
| Refinement Statistics: |  |  |
| Resolution (Å) | 41.79 - 2.29 | 41.2 - 1.8 |
| No. Reflections | 7549 | 12785 |
| $R_{\text{work}} / R_{\text{free}}$ | 0.209/0.245 | 0.215/0.224 |
| No. Atoms | 1123 | 993 |
| Protein | 1057 | 940 |
| Water | 66 | 53 |
| Average B, all atoms (Å<sup>2</sup>) | 54.3 | 50 |
| R.M.S. Deviations: |  |  |
| Bond lengths (Å) | 0.002 | 0.007 |
| Bond angles (°) | 0.433 | 0.965 |
| Protein Data Bank Entry | 6GUV | 6GUW |

\* Values in parenthesis are for the highest-resolution shell.

### 2.4. Structure solution and refinement

The structure of the mouse PATZ1 BTB was solved by molecular replacement with PHASER (Mccoy et al., 2007) using the PLZF BTB structure (PDB entry 1BUO) as search template. The identified solution was then subjected to rounds of manual rebuilding with the program COOT (Emsley et al., 2010) and refinement with PHENIX (Liebschner et al., 2019) to give final R_work_ and R_free_ of respectively 20.9% and 24.5%. The final structure was validated with MOLPROBITY (Williams et al., 2018) and deposited with the structure factors in the protein data bank under accession number 6GUV.

The structure of the zebrafish PATZ1 BTB was solved by molecular replacement with PHASER using the mouse PATZ1 BTB structure (PDB entry 6GUV) as search template. The identified solution was then subjected to rounds of manual rebuilding with the program COOT and refinement with PHENIX to give final R_work_ and R_free_ of respectively 21.5% and 22.4%. The final structure was validated with MOLPROBITY and deposited with the structure factors in the protein data bank under accession number 6GUW.

Detailed X-ray data refinement statistics are given in Table 1.

### 2.5. Sequence and Structure Analysis

Sequences of PATZ1 protein of different organisms were retrieved from NCBI’s reference sequence (RefSeq) database (Pruitt *et al.*, 2007). Except for *Homo sapiens* (NP_114440.1) and *Mus musculus* (NP_001240620.1) for mammals, one organism only is chosen for each group of different species of vertebrates: *Danio rerio* (XP_009300883.1) for fish, *Xenopus laevis* (XP_018117120.1) for amphibians, *Thamnophis sirtalis* (XP_013922905.1) for reptiles and *Parus major* (XP_015499085.1) for birds. Sequence limits are determined based on the annotations of the BTB domain in the UniProt database (Bateman *et al.*, 2019). The 49 members of the human ZBTB protein family were retrieved from Swiss-Prot and a multiple sequence alignment was obtained by PROMALS3D online tool (Pei et al., 2008). This alignment incorporates the structural information from the available PDB structures of these proteins. Secondary structure nomenclature refers to Stogios et al. 2005 (Fig. 1B and 2A). ZBTB4 was excluded from this alignment because of its N-terminal serine-rich repetitive insertions which are beyond the scope of this description. Shades were added according to the percentage of similarity for every position in the alignment visualized in Jalview-2 (Waterhouse et al., 2009) that generated a consensus sequence. A list of residues involved in homodimer interaction interfaces was obtained by PDBePISA (Krissinel & Henrick, 2007) and graphically rendered on the protein structure with VMD (Humphrey et al., 1996). Intra- and inter-chain interactions were retrieved by PIC-Protein Interactions Calculator (Tina et al., 2007) and VMD Salt Bridges plug-in and Timeline. Structural alignments were calculated on the MultiProt server (Shatsky et al., 2004).

### 2.6. Modelling

The central loop of the mammalian PATZ1 BTB structure (residues 83-113) was modelled as a monomer by using the PRIMO suite (Hatherley et al., 2016) based on the MODELLER program (Sali & Blundell, 1993). Structural information from MEME motif analysis of the loop sequence was added to the template (Bailey & Elkan, 1994). The obtained model was aligned to the deposited structure (6GUV) in PyMOL (Schrodinger, 2015). The coordinates of the loop model were added to the crystal structure and fragments were joined using the VMD AutoPSF plugin. SymmDock (Schneidman-Duhovny et al., 2005) was used to reconstruct the dimer conformation.

The stability of the new structure was tested by molecular dynamics (MD) simulations on NAMD (Phillips et al., 2005). The protein structure model was centered in a solvent box built according to the protein size and padded with at least a 10 Å layer of water in every direction. The solvent was modelled explicitly using TIP3W water molecules. 0.15 M KCl was added to ionize the solvent. The MD simulation was performed using the Charmm27 force field (Brooks et al., 2009) on the NAMD program. Periodic boundary conditions were applied in which long-range electrostatic interactions were treated using the particle-mesh-Ewald method (Darden et al., 1999) and the cut-off distance was set to 12 Å. All simulations were run in duplicates at constant temperature of 310 K for at least 200 ns.

In one of the MD simulations of the human PATZ1 BTB, the flexible loop region in one of the monomers formed an extra β-strand structure. The model was further refined by once more duplicating this monomer into the dimer form. This refinement was processed by taking a frame from the run where the root mean square deviation (RMSD) with the initial structure, excluding the loop, was minimal (3.04 Å) and by mirroring the information from the monomer with the formed secondary structures to the other monomer with M-ZDOCK (Pierce et al., 2005), recreating the dimer by symmetry. The stability of the new structure was confirmed by additional MD simulations (twice, 200 ns each). ModLoop (Fiser & Sali, 2003) was used to model the coordinates of the missing residues (70-76) in the zebrafish PATZ1 BTB structure (6GUW) and the modelled structure stability was assessed by MD as before.

## 3. Results and discussion

### 3.1. Structural features of the murine and zebrafish PATZ1 BTB domains

Here we report the crystal structure of the BTB domain of murine and zebrafish PATZ1 (Fig. 2). Similarly to other members of the ZBTB family, both PATZ1 BTB crystal structures reveal a strand-exchange homodimer, organized as a core fold BTB domain, preceded by an N-terminal extension that interacts with the partner chain in the dimer. The characteristic secondary structures of the dimerization interfaces (β1-α1; A1-A2; β2-A5) are conserved. Size exclusion chromatography data for both murine and zebrafish BTB domains suggests that the homodimeric complex is the most abundant oligomerization state found in solution (data not shown). The murine PATZ1 BTB domain protein was expressed from a construct encoding amino acids 12-166 preceded at the N-terminus by 20 amino acids comprising the His-tag and an HRV-3C protease digestion site (Fig. 2A-B). The last 10 of these amino acids (ALEVLFQGPG) are visible in the structure and fold into a β-strand (β0), anti-parallel to the first N-terminal PATZ1 β-strand (β1). Superposition of this structure with that of BCL6 BTB in complex with co-repressor peptides (PDB entries 1R29, 3BIM) suggests that these extra residues structurally mimic the β-strand forming residues from both SMRT and BCOR (Supp. Fig. 1) although their amino acid sequence is not conserved. Attempts to crystallize the mouse PATZ1 BTB construct following cleavage of the His-tag failed, suggesting that the extra N-terminal amino acids likely aid the crystallization process in this case. Thus, the groove of the BTB domain, that binds co-repressor peptides in BCL6, is promiscuous, potentially accommodating various ligands that do not necessarily share a conserved sequence.

**Figure 2.**
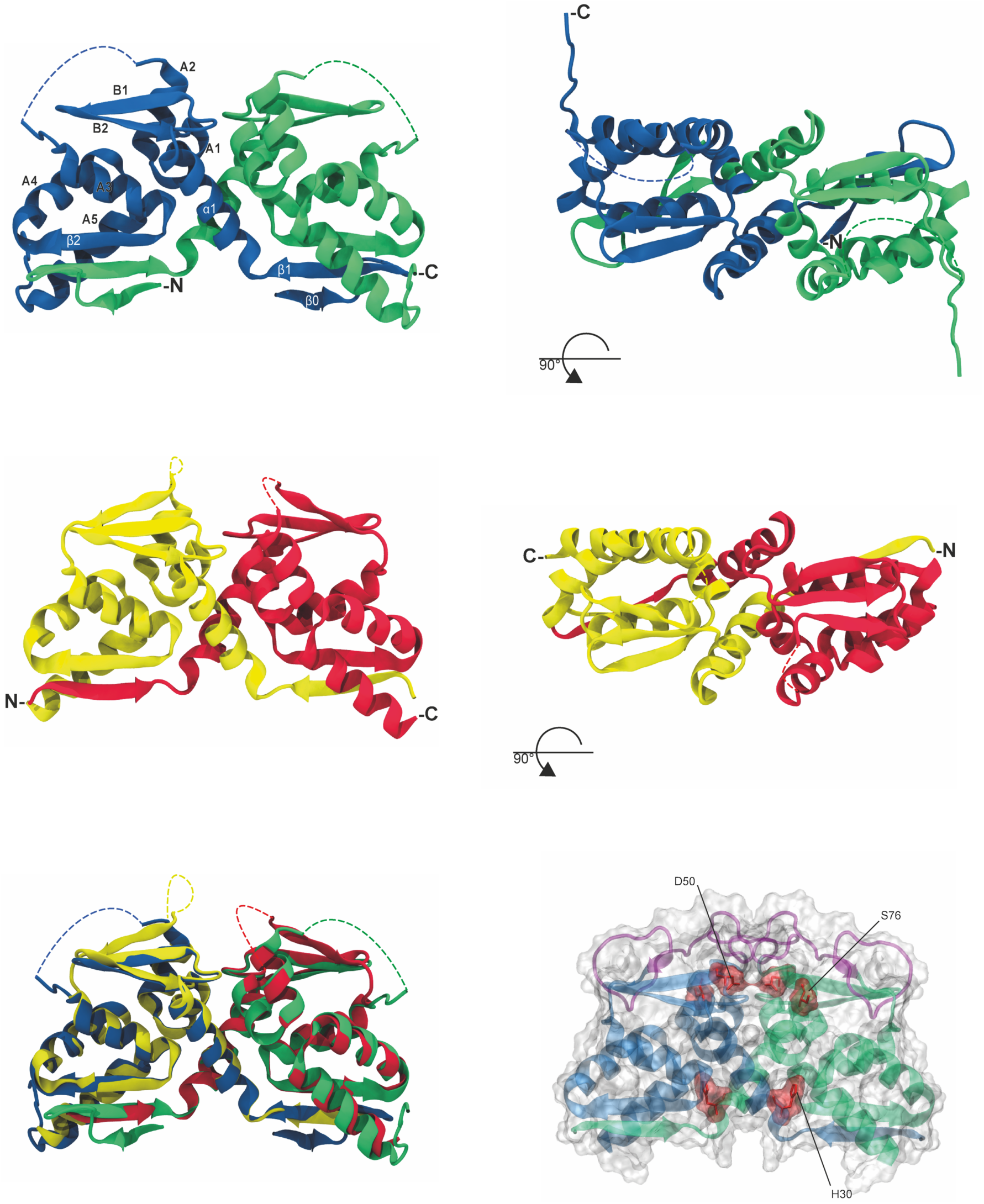
Structure of the PATZ1 BTB Domain. **(A)** Crystal structure of the BTB domain of the mouse PATZ1 protein (PDB entry 6GUV) in cartoon representation (front view). The crystallographic asymmetric unit contains a PATZ1 BTB domain dimer colored blue and green. Secondary structures are indicated in capital and Greek letters. **(B)** Top view of the mouse PATZ1 BTB structure. N- and C-termini are indicated on one monomer. The coordinates of 31 residues in a central region, unique to mammalian PATZ1 BTB domains, could not be assigned (indicated by dotted loop). **(C)** Crystal structure of the BTB domain of the zebrafish PATZ1 protein (PDB entry 6GUW) colored in yellow and red (front view). **(D)** Top view of the zebrafish PATZ1 BTB structure. The coordinates of 7 residues in the zebrafish PATZ1 BTB domains could not be assigned (indicated by dotted loop). **(E)** Comparison of the mouse (blue and green) and zebrafish (yellow and red) PATZ1 BTB domains (RMSD 1.02 Å). **(F)** A space filling representation of the mouse PATZ1 BTB domain structure. The predicted structure (in purple) of the central region was generated by homology modelling followed by conformation equilibration using MD simulations. Note that the model structure contains a predicted short β-strand. Highlighted in red, the three conserved degron residues, annotated at the bottom of Fig. 1B and predicted to play a role in BTB dimer degradation are buried. Numbering refers to the residues in the crystal structure.

Mammalian PATZ1 (ZBTB19) protein is predicted to contain a 31 amino acid long A2/B3 loop (residues 75-105) partially conserved in cKrox (ZBTB15), which sets PATZ1 apart from the other ZBTB family members (Fig. 1A and B). This large loop replaces a shorter amino acid stretch that in other ZBTB family proteins forms a β-strand (B3), described in detail by Stogios et al. (Stogios et al., 2005). This loop is glycine and alanine rich and is predicted to be partially disordered, however a short β-strand is predicted for the last 6 residues. The single difference between the human and mouse PATZ1 BTB domains (T91A) is found within this large loop. In the crystal structure, no density could be assigned to residues belonging to the A2/B3 loop, suggesting that these amino acids are at least partially disordered or flexible.

Interestingly, seven-amino acids stretch from the C-terminus of an adjacent molecule in the crystal unit cells extends into the region normally occupied by the B3 β-strand in other ZBTB proteins. Presumably this fragment mimics B3 and stabilizes the β-sheet formed by B1 and B2 in the mouse PATZ1 BTB domain. To test this hypothesis, we crystallized a construct lacking the last seven amino acids in the C-terminus of PATZ1 BTB. Crystals for this protein diffracted poorly and to lower resolution (3.4 Å), showing a different packing (data not shown). This suggests that the seven amino acids were in fact important for stabilizing the B1-B2 β-sheet and for crystal packing, yet their absence did not encourage the folding of the A2/B3 loop into a β-strand (Supp. Fig. 2).

Sequence alignment shows that the length of the A2/B3 loop is conserved in all PATZ1 orthologues but is conspicuously absent in fish and amphibians (Fig. 1A). The mammalian A2/B3 loop could be an evolutionarily acquired insertion sequence encoding an intrinsically disordered loop (IDL) (Fukuchi et al., 2006). To study the structure of this region in detail, we solved the structure of the zebrafish PATZ1 BTB domain (residues 1-135) to a resolution of 1.8 Å (Fig. 2C-D). When excluding the central loop, the murine and zebrafish sequences share 83.7% identity whilst the structures can be superimposed with a root mean square deviation (RMSD) of 0.62 Å (Fig. 2E). The zebrafish BTB domain also shares the same quaternary structure of the mouse BTB, a strand-exchange homodimer. The zebrafish PATZ1 BTB domain is however structurally more similar to other ZBTB proteins than the mammalian PATZ1 BTB, which stands alone as an outlier. The zebrafish PATZ1 BTB domain also has a seven amino acid long sequence between A2 and B3 for which no discernible electron density could be found. This loop was modelled using ModLoop (Fiser et al., 2000, Fiser & Sali, 2003) (Fig. 3D).

**Figure 3.**
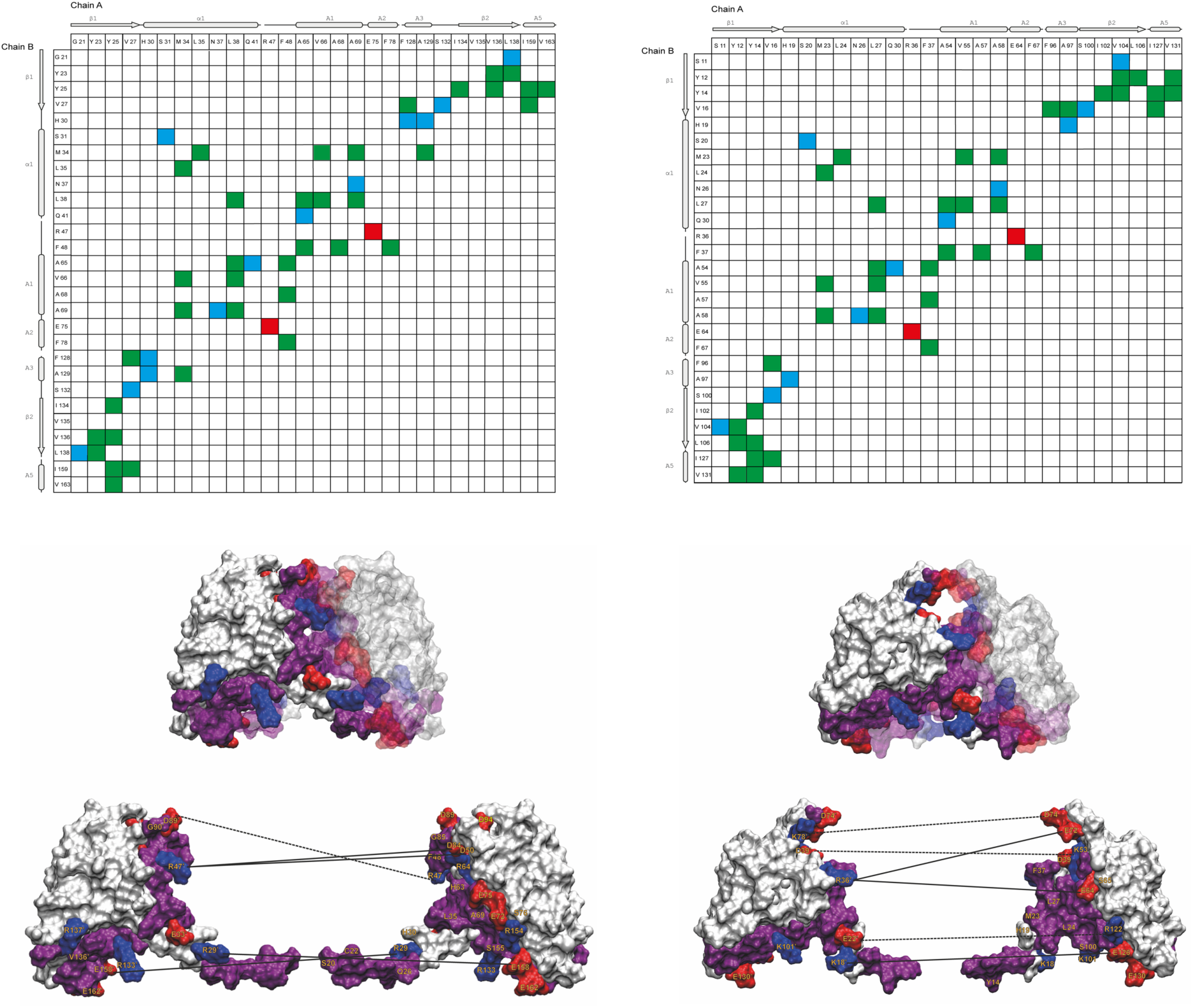
Identification of residues involved in homodimer interaction interfaces. Contact maps of interchain residue interactions of the mouse **(A)** and zebrafish **(B)** PATZ1 BTB crystal structures. Hydrophobic interactions (green), hydrogen (blue) and ionic bonds (red) are indicated. The relevant elements of the secondary structures are shown for orientation. A highly charged dimerization interface mediates PATZ1 BTB homodimerization. **(C)** A split homodimer view in surface representation and completed with the modelled loop, highlights residues involved in the interaction interface; positively (blue) and negatively (red) charged residues are annotated, neutral residues are shown in purple. Interchain salt bridges that persist above the threshold are indicated by straight lines and those that do not persist by dotted lines. **(D)** The zebrafish PATZ1 dimerization interface is also shown in split homodimer view for comparison. Numbering refers to the crystal structures files.

The human and mouse PATZ1 BTB domains have a single amino acid difference (T91A). To detail the possible structure and dynamic behavior of the human PATZ1 BTB A2/B3 loop, we performed molecular modelling and dynamics simulations (MD). MD simulations lasting 200 ns indicated that while the overall dimer forms a stable structure, the A2/B3 loop region is uniquely flexible. The simulations also suggest that a new β-strand could form within this loop (Fig. 2F and 3C) and that at least 10 amino acids within the modelled loop contribute to the homodimerization interface (representing 17.5% of the total interface of 57 amino acids). Interestingly, the β-strand that is generated in the simulations contains the threonine residue which is the only amino acid that is different between the human and mouse BTB domains. The flexibility of this loop in MD simulations indicates that the loop may be dynamically modulating homodimer stability. Other BTB domains, like LRF and MIZ1 also contain flexible loops in this region (Stogios et al., 2007, Stead et al., 2007). In the case of MIZ1, the A2/B3 region mediates tetramerization of its BTB domain, whilst using size exclusion chromatography and small angle X-ray scattering experiments (SAXS) we found no evidence for such oligomerization in mouse or zebrafish PATZ1 (data not shown).

### 3.2. A highly charged and dynamic surface contributes to the stable homodimerization interface of the PATZ1 BTB domain

The mouse and zebrafish PATZ1 BTB domain dimerization interfaces are very similar. Using the PIC tool (Tina et al., 2007) we determined that their crystal structures contain a single structurally corresponding salt bridge (ARG47-GLU75 in mouse and ARG36-GLU64 in zebrafish; residue numbering refers to the crystal structures), in addition to the residues engaged in inter-chain hydrophobic interactions and hydrogen bonds (Fig. 3 A-B). The interfaces retrieved from PDBePISA (Krissinel & Henrick, 2007), contain four basic and five acidic residues (murine) and four basic and three acidic residues (zebrafish) (Table 2).

**Table 2.**
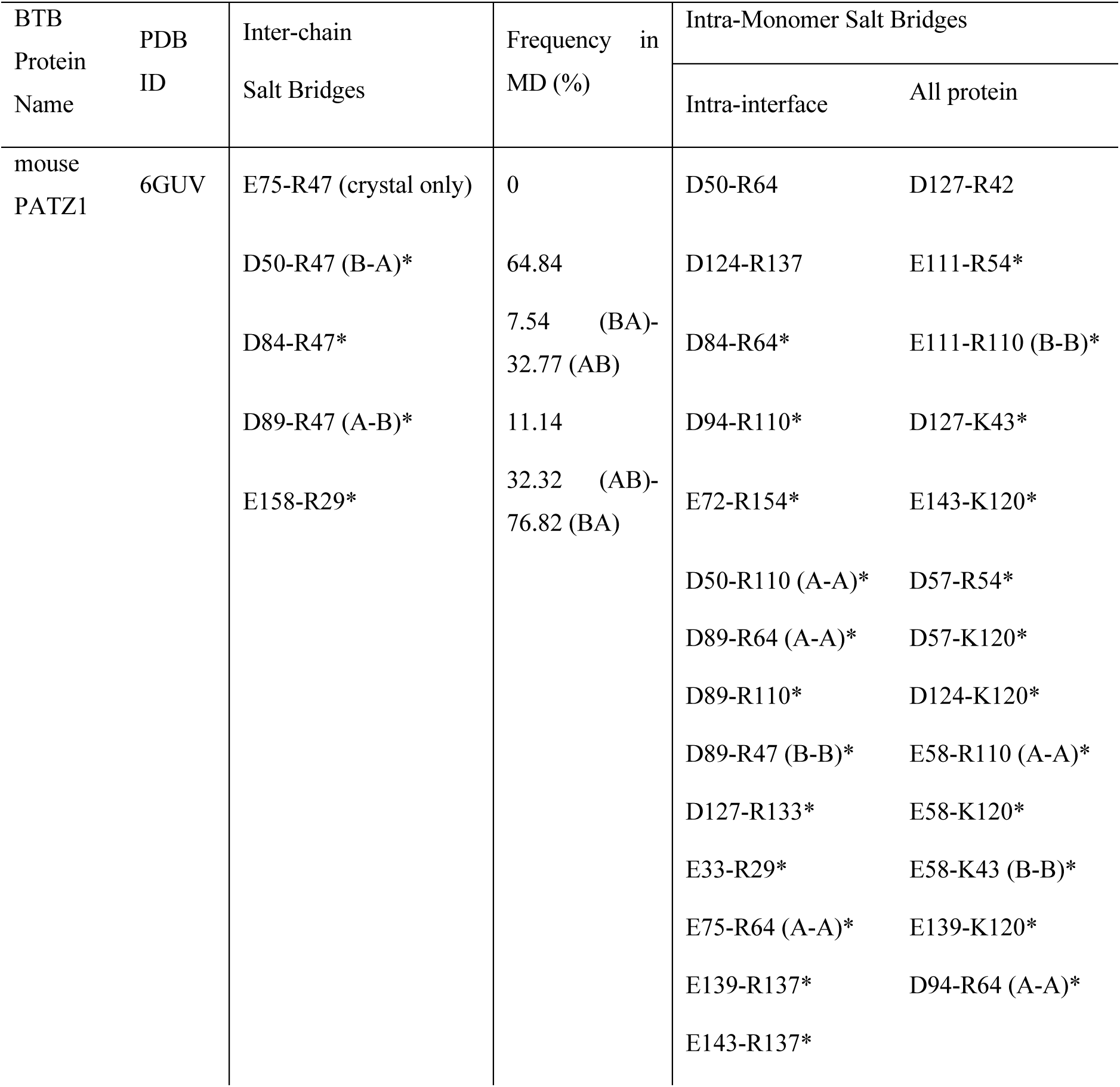
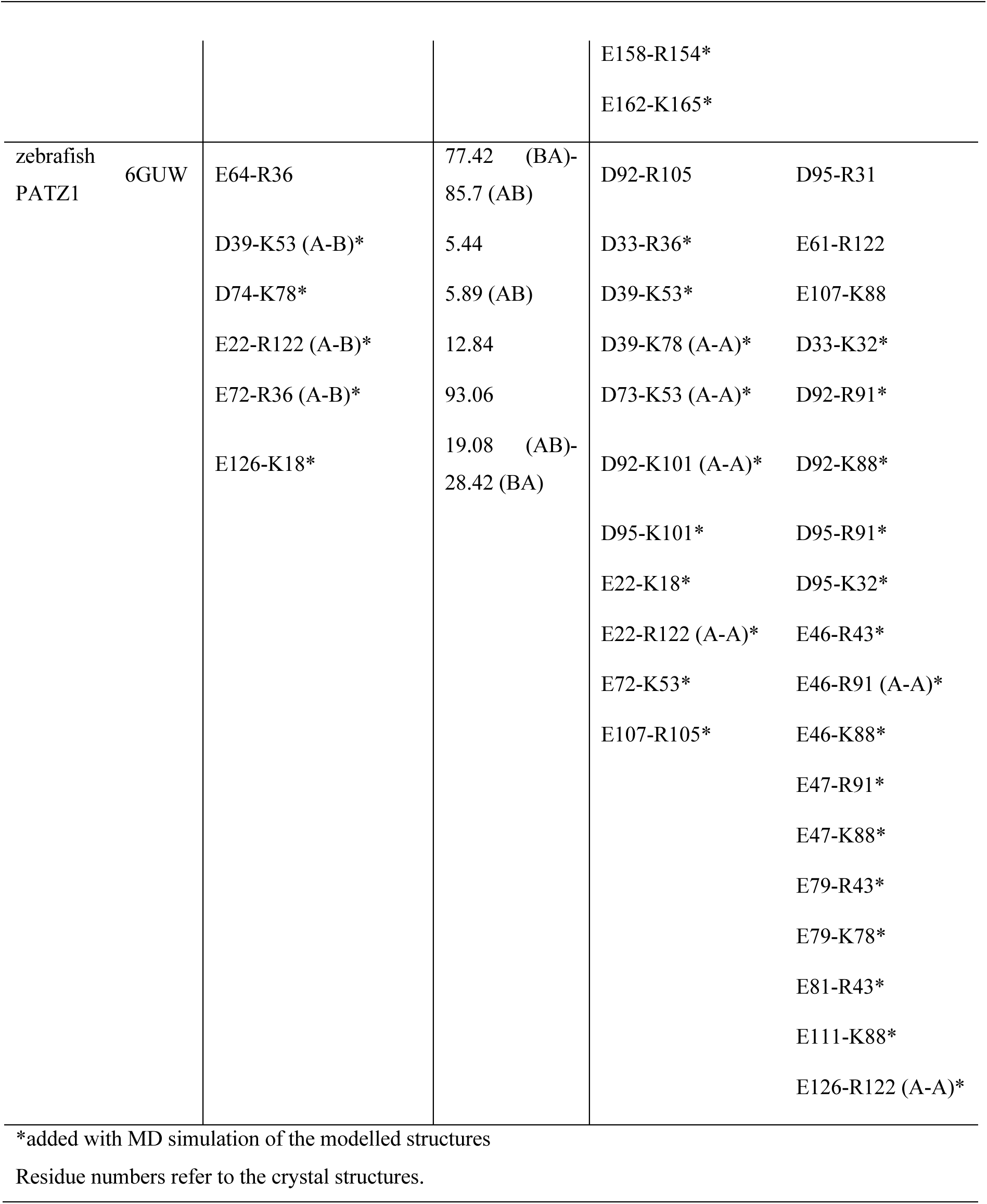
Inter-chain salt bridges in the dimerization interface of BTB domain of PATZ1

In order to understand the dynamics of these interfaces, we assessed the number of contact-forming residues in the energy minimized modelled BTB domains (Fig. 2F and Fig. 3C-D). Using the VMD tools, we found a dramatic increase of charged residues (mostly negative) that participate in interface contacts (Table 2 asterisks). To understand the stability of these contacts, we assessed those that persist above a threshold value (15%) during the lifetime of the MD simulation (Fig. 3 C-D). During the MD, the A2/B3 loop region significantly contributes to the interface in both models, resulting in flexibility of salt bridges that form between a single charged amino acid from one monomer with multiple opposite charged amino acids from the opposite monomer. While ARG47 of mouse PATZ1 BTB, is engaged only with GLU75 in the crystal structure, MD shows that it can contact a broader number of charged residues, including those from the flexible loop (ASP50, ASP84, ASP89). We also find that two of the three degron residues (annotated in Fig. 1B and Fig. 2F) which are predicted to play a role in BTB heterodimer degradation, participate in the interaction interface of both BTB homodimer structures.

### 3.3. Co-repressor binding modalities are not conserved in different BTB domains

BTB domains have been shown to interact with co-repressor proteins NCOR1, SMRT and BCOR (Huynh & Bardwell, 1998, Wong & Privalsky, 1998, Melnick *et al.*, 2002, Huynh *et al.*, 2000). Co-repressor binding to the BCL6 BTB domain requires dimerization because the interaction interface (lateral groove) is formed by residues on both monomers. Twenty-three residues from each BCL6 monomer contribute to this interface (Ahmad *et al.*, 2003) (Fig 4). Additionally, four residues (L19S; N23H; L25S/R26L) when mutated, interfere with co-repressor binding by preventing homodimerization of the BTB domain (Huynh & Bardwell, 1998, Ghetu *et al.*, 2008, Granadino-Roldan *et al.*, 2014). The PATZ1 BTB domain has also been shown to bind to NCOR1, suggesting that a similar lateral groove may be mediating this interaction (Bilic et al., 2006). In fact, when the residues corresponding to BCL6 L19S; N23H; L25S/R26L were mutated in the PATZ1 BTB domain (L27S, Q33S, R34L), it also failed to bind NCOR1 (Bilic *et al.*, 2006).

**Figure 4.**
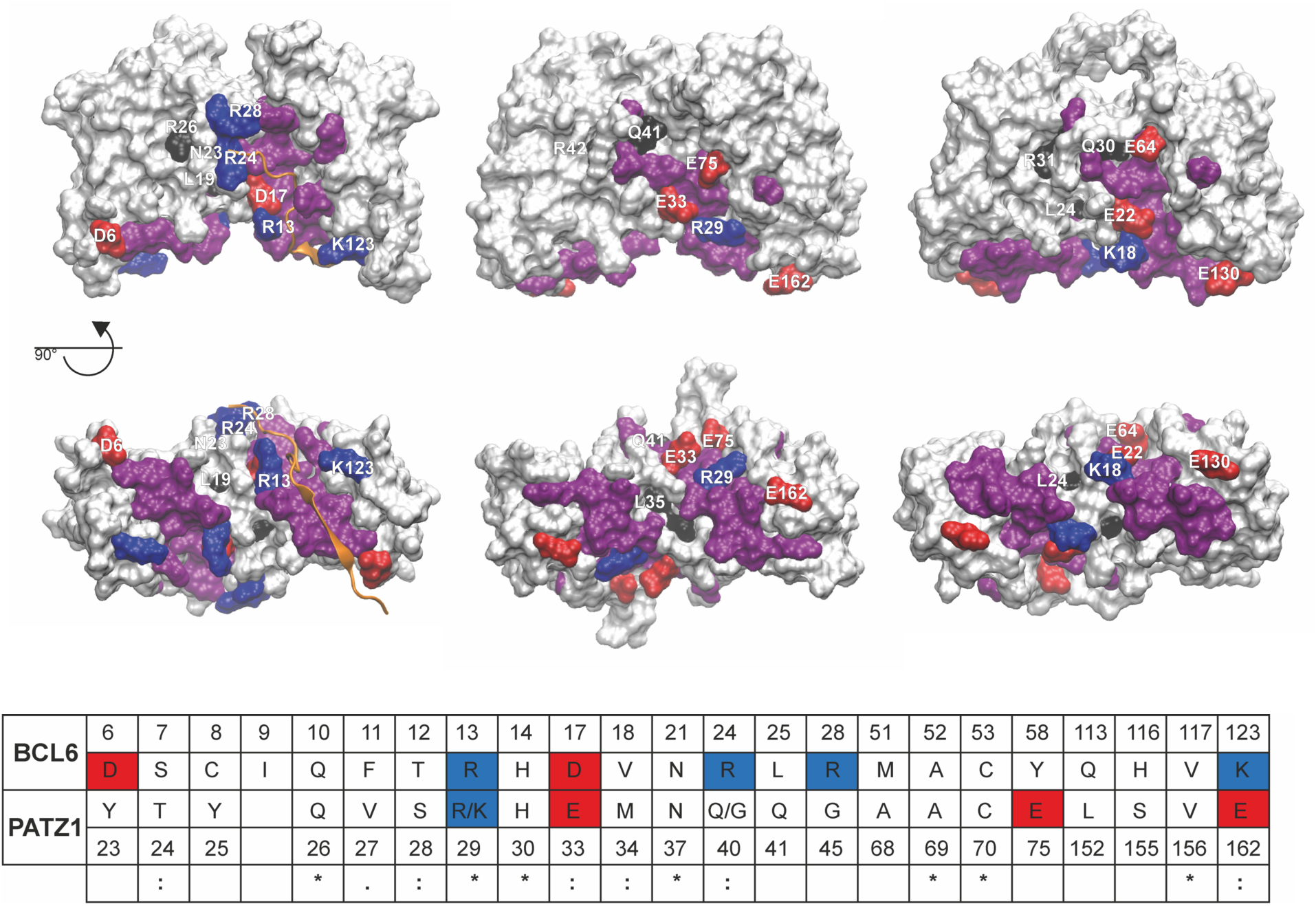
Comparison of the lateral grooves of PATZ1 and BCL6 BTB domains. The structures of the BCL6 (PDB entry 1R2B) **(A)**, and the energy minimized modelled mouse **(B)** and zebrafish **(C)** PATZ1 BTB domains are shown in surface representation in front and bottom view. The surface area of the residues in the lateral groove of BCL6 BTB domain is buried upon formation of BCL6-SMRT complex (Ahmad, 2003). The SMRT peptide binding the BCL6 lateral groove is shown in cartoon representation in orange. All residues in the lateral groove are labelled for one monomer. Colors indicate residue type: positive charged (blue), negative charged (red). All other residues in this region are colored in purple. The position of the mutations that in BCL6 and in PATZ1 affect the binding with the co-repressor peptides is indicated in black (references in the text). A sequence alignment of BCL6 and PATZ1 BTB residues located in the lateral groove region is shown **(D)**. Residues are numbered according to the BCL6 and mouse PATZ1 structure files with the charged residues colored as in (A)-(C). Apart from position 29 and 40 where the two alternatives are indicated, mouse and zebrafish PATZ1 contain the same residues in these structurally corresponding positions. Although residue conservation for BCL6 and PATZ1 in this region is low, SMRT/NCOR peptides are predicted to bind the BTB domain of PATZ1 in the same region.

Even though BCL6 and PATZ1 are structurally very similar, their corresponding co-repressor binding interface sequences are not conserved (Fig. 4 D). To examine the structural similarity between the PATZ1 and BCL6 BTB domains, we calculated the RMSD (1.56 Å) between individual monomers. The structural similarity between BCL6 and PATZ1 was more evident when the flexible PATZ1 loop was excluded (RMSD of 1.23 Å). Comparison of the surface charge distribution of the two proteins indicates major differences (Fig. 5). Specifically, the BCL6 lateral groove contains a high density of positively charged amino acids that interact with the co-repressors (the interaction with SMRT and BCOR are shown in Fig. 5). Surprisingly, the surface of PATZ1, corresponding to the BCL6 lateral groove did not contain as many basic residues (Fig. 4 D and Fig. 5). In fact, this region of mouse and zebrafish PATZ1 are highly conserved (91% identical) and contain more acidic amino acids.

**Figure 5.**
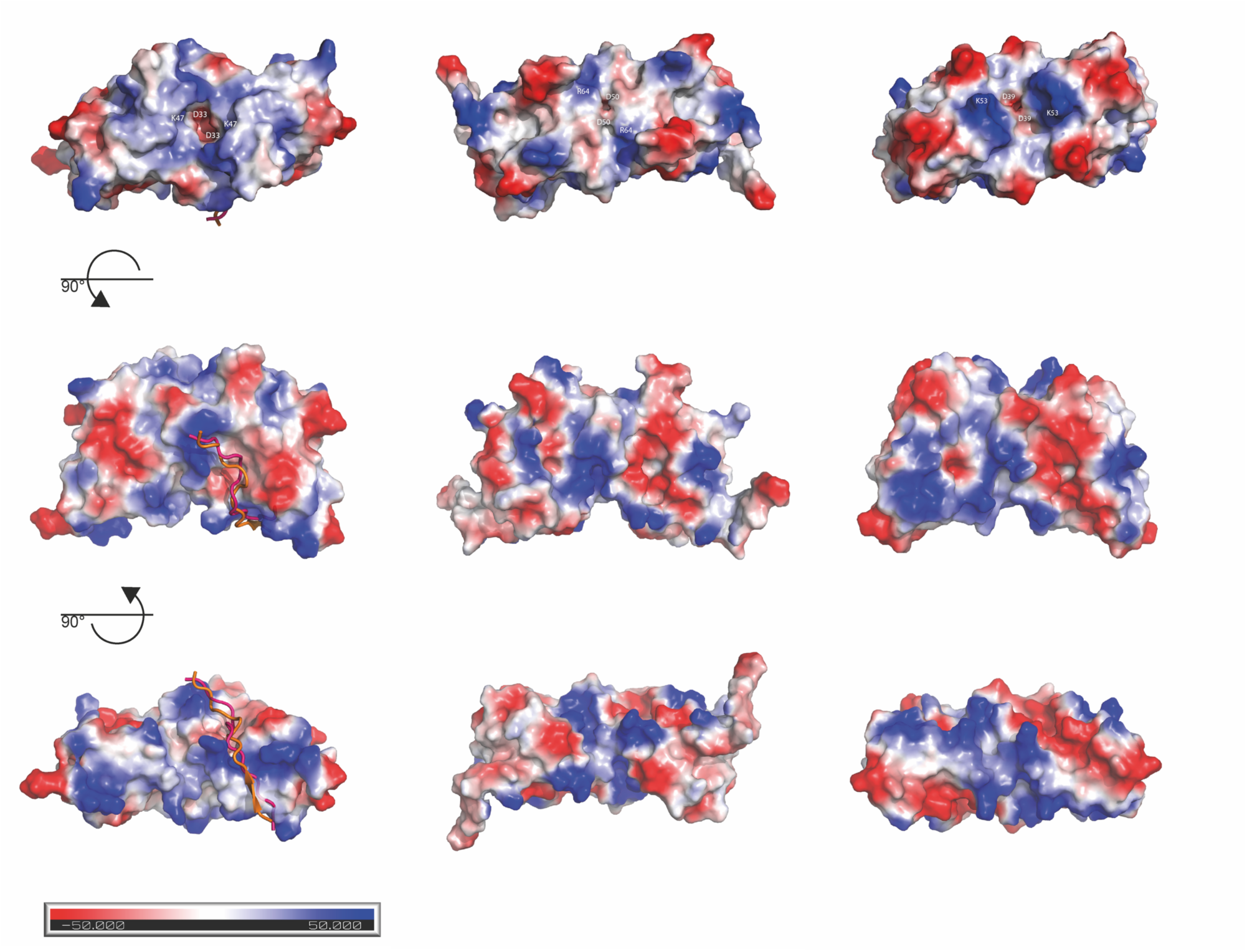
Comparison of the electrostatic surface potentials of PATZ1 and BCL6 BTB domains. From the central front view, two 90° rotations show top and bottom views of the electrostatic surface potential of the BTB domain of: **(A)** BCL6 bound to the SMRT BBD co-repressor peptide (cartoon representation magenta) and the overlapped BCOR (in orange) (PDB entries 1R2B and 3BIM); **(B)** murine PATZ1 (6GUV); **(C)** zebrafish PATZ1 (6GUW). In the top view two conserved charged residues (negative and positive) that characterize the BTB domain charged pocket are indicated. Residues are numbered according to the BCL6 and PATZ1 structure files. The colors range from red (negative potential) to blue (positive potential) with white near neutral.

The presence of alternatively charged residues in the lateral groove of PATZ1 may indicate that its interaction with co-repressors may be through a different mode compared to BCL6. In this regard, the lateral groove of PATZ1 BTB domain is more similar to that of LRF compared to BCL6 (Stogios *et al.*, 2007). We also find that residue D50 in the structure of mouse PATZ1, corresponding to D39 of zebrafish and to D33 of BCL6 (Fig. 5), which is absolutely conserved in all ZBTB proteins and happens to be the previously mentioned second degron residue Fig. 1, is a part of the charged pocket that is conserved between BCL6, LRF and PLZF (Stogios *et al.*, 2007). This charged pocket that is formed by the participation of D50 residues from both monomers has been suggested as an alternative region for ligand binding (Fig. 5).

### 3.4. Discussion

A role for PATZ1 has been shown in various malignancies such as thyroid and testicular cancer (Fedele *et al.*, 2008, Valentino, Palmieri, Vitiello, Pierantoni, *et al.*, 2013, Chiappetta *et al.*, 2015, Vitiello *et al.*, 2016, Fedele *et al.*, 2017, Monaco *et al.*, 2018). The interaction between PATZ1 and the tumor suppressor p53 (Valentino, Palmieri, Vitiello, Pierantoni, *et al.*, 2013, Valentino, Palmieri, Vitiello, Simeone, *et al.*, 2013, Chiappetta *et al.*, 2015, Keskin *et al.*, 2015) mediated by a motif in the zinc finger DNA binding domain rather than the BTB domain also links this protein to cancer. Chromosome 22 specific inversions that translocate the transcription factor EWSR1 with PATZ1 have been observed in various sarcomas. While EWSR1 translocates with ETS family proteins in Ewing’s sarcoma, it potentially encodes two fusion proteins EWSR1-PATZ1 and PATZ1-EWSR1, the latter of which will contain the BTB domain (Siegfried *et al.*, 2019, Bridge *et al.*, 2019, Chougule *et al.*, 2019, Sankar & Lessnick, 2011, Mastrangelo *et al.*, 2000, Im *et al.*, 2000). The dimerization and co-repressor interaction properties of PATZ1 identified in this study may shed light to the mechanism of these sarcomas.

The gene targets of PATZ1 have not been extensively identified. ChIP-Seq and RNASeq experiments have identified 187 putative targets (Encode Project Consortium 2012, Keskin *et al.*, 2015). Of these targets roughly half were upregulated, and half downregulated in the absence of Patz1 expression. How many of these genes are direct targets and how many require the BTB domain for regulation is not known. The current study only highlights structural motifs that likely play a role in the interaction of the PATZ1 BTB domain with co-repressor proteins. Yet other interactions may be involved in the potential role of PATZ1 in gene upregulation.

Our present study identifies important structural features of the PATZ1 BTB domain. One unique feature of BTB domains is their ability to form homodimers as well as heterodimers as a result of their close structural homology. Dimer formation is necessary for interaction with co-repressor proteins for both PATZ1 and BCL6. A lateral groove that BCL6 uses to bind co-repressors is structurally conserved in PATZ1 (Fig. 4). However, the discrepancy of charged amino acids in this groove (Fig. 5) may indicate altered binding modalities and/or affinities to co-repressors. In fact a structurally conserved charged pocket has previously been hypothesized to be involved in ligand interaction (Stogios *et al.*, 2007). While in BCL6, LRF and PLZF structures, this charged pocket is surface exposed, the PATZ1 A2/B3 loop could dynamically gate this site, potentially regulating ligand interaction.

Another feature that is common between PATZ1 and BCL6 is their BTB domain mediated localization to nuclear speckles (Huynh *et al.*, 2000, Fedele *et al.*, 2000, Franco *et al.*, 2016). While PATZ1 interacts with the nuclear speckle resident ubiquitin ligase RNF4, whether potential post-translational modifications due to this interaction affect its stability is not known (Pero *et al.*, 2002). A recent study identified three BTB domain degron residues that are surface exposed preferentially in heterodimers (Mena *et al.*, 2018). Targeting of these degrons by ubiquitin ligases mediates proteasome dependent degradation of heterodimers over homodimers. It is not known if this is a generalizable feature of BTB domains. Consistent with the stability of homodimers, in the current study, we find that PATZ1 buries two out of three of these degron residues in the protein globular structure (Fig. 2F).

The MD simulation shows that alternative contacts are possible for several charged residues at the homodimer interface (Table 2). Because a single charged residue can contact more than one opposite charged residue (Fig. 3), the dimer interface may be more resilient to the disruption of single contacts. Energetically, the presence of alternative salt bridges may be necessary to accommodate the flexibility of the central loop whilst retaining the stability of the dimerization interface. Nevertheless, there is an exceptional number of unpaired charged surface residues in the murine PATZ1 BTB domain. Some proteins, eg. calmodulin, with large net surface charges are known to modulate their environment by redistributing nonspecific ions in the surrounding medium (Aykut et al., 2013). Such modes of action are utilized to shift the population of the available conformational states leading to fine-tuned functions.

BTB domains are attractive targets of anti-cancer compounds. Compounds that prevent homodimerization or result in degradation of BCL6 (Kerres et al., 2017) have been shown to have highly effective cytotoxic activity in B lymphomas. Because the BTB domains of ZBTB family proteins all share the same fold, compound specificity requires targeting unique features. The residues in the A2/B3 loop of PATZ1, unique among the ZBTB proteins, are potentially a specific target for this protein (Fig. 1). The structure of the PATZ1 BTB domain reported in this study will aid in the development of therapeutics for those human malignancies that involve PATZ1 and the testing of the specificity of compounds targeting other BTB domains.

## Supporting information

Supporting informaton

## Acknowledgements

This work was funded by TUBITAK 1001 grant 118Z015 and The Royal Society, Newton International Exchanges Grant NI140172. S.P. was supported by a TUBITAK BIDEB scholarship. S.P. and A.A. performed experiments and solved the crystal structures; S.P. and C.A. performed the MD and S.P., B.E. and E.M. supervised the overall study and wrote the manuscript. We thank Prof. Dr. Stefan H. Fuss, Prof. Dr. Zehra Sayers, Prof. Dr. Wilfried Ellmeier for helpful comments for the preparation of the manuscript.

