## Supporting informaton for "Structural Analysis of the PATZ1 BTB domain homodimer"

### Supporting information

**Figure S1** Superposition of the BCOR bound BCL6 BTB domain monomer structure with that of the mouse PATZ1 BTB domain dimer. His-tag residues form an N-terminal β-strand of the PATZ1 BTB domain construct to mimic the β-sheet interaction of co-repressors with BCL6. PATZ1 homodimer is shown in green and blue, BCL6 monomer is shown in violet and BCOR co-repressor peptide is shown in orange.

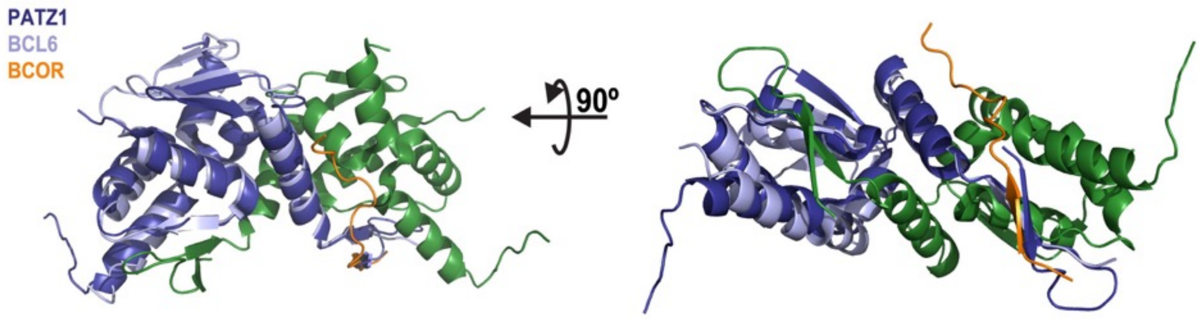


**Figure S2** A C-terminal sequence from an adjacent molecule mimics the B3 B-strand to stabilize the β-sheet formed by B1 and B2 strands in the mouse PATZ1 BTB domain. Two PATZ1 BTB monomers forming a homodimer in the crystal unit are shown in green and blue respectively. A third monomer from an adjacent molecule in the crystal (shown in magenta) contributes to the β-sheet formed by B1 and B2.

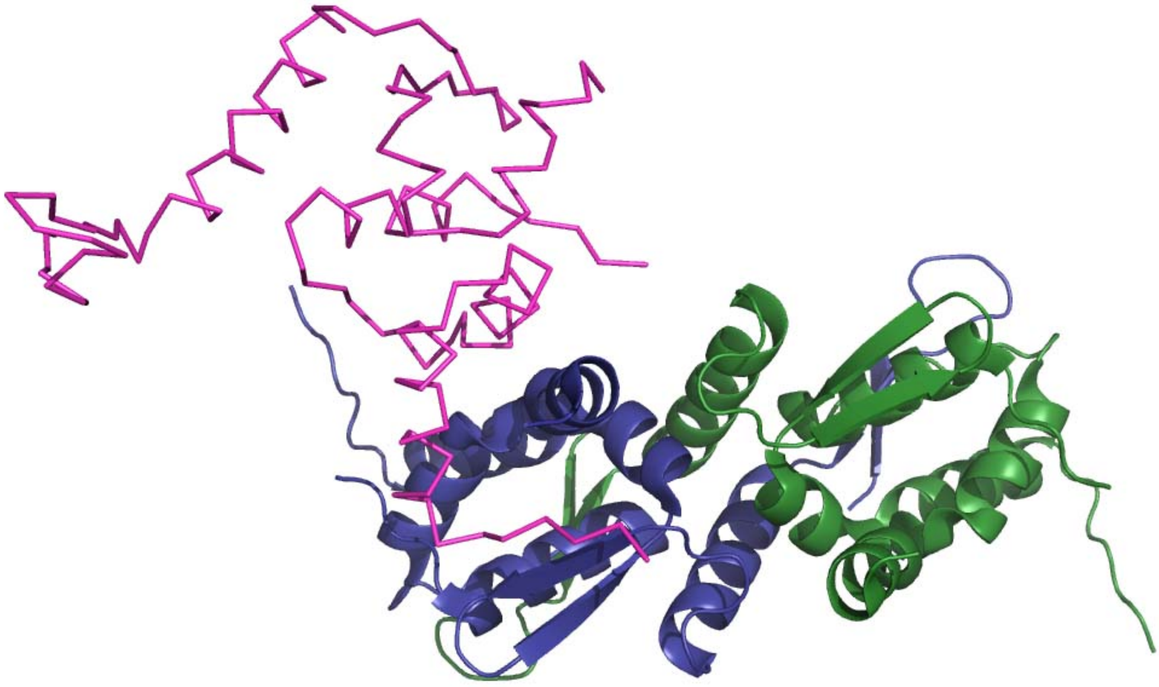
